# dnaSORA - A Unified Diffusion Transformer for DNA point clouds

**DOI:** 10.1101/2025.01.27.633223

**Authors:** Oleksandr Koreniuk, eMalick G. Njie

## Abstract

The relatively obscure Hawaiian experiment collapses diverse phenotypes, including nearly all human genetic diseases to a singular Gaussian-like point cloud feature, structuring unstructured information. The uniformity of the feature space provides a straightforward way for AI models to learn all three billion tokens for reading the human genome as a first language. We propose a diffusion transformer, dnaSORA, for learning these features. dnaSORA has generative capacity similar to Stable Diffusion but for DNA point clouds. The model’s architecture is novel because it is unified; thus, it also functions as a discriminator that uses a frozen latent representation for classification. dnaSORA transfer learns from synthetic data emulating real genome point clouds to classify misrepresented tokens in *C. elegans* Hawaiian data at state-of-the-art 0.3 Mb resolution. Pre-training large genome models typically requires expensive and difficult-to-obtain genomes. However, our solution provides nearly unlimited synthetic training data at negligible compute costs. Inference for new token assignments (e.g., new diseases) requires genomes from several dozen rather than thousands of individuals. These efficiencies, combined with state-of-the-art resolution, provide a pathway for rapid, massive scaling of token annotation of the entire human genome at orders of magnitude below expected costs.

## Non-expert description

What is possible if one can understand the human genome as a first language? What does understanding something as a first language even mean? And how would you go about obtaining such understanding? We at Ecotone have been exploring these questions. And have achieved significant progress in getting answers. For instance, such understanding will result in improved development of CRISPR medicines for thousands of rare genetic diseases.

Think of reading the human genome like trying to understand a mysterious ancient language with 3 billion words - both humans and AI have struggled to make sense of it. We’ve combined a relatively obscure genetic mapping technique called the Hawaiian experiment with modern AI technology. We found that despite the vast complexity of human genetic diseases, they all create similar, predictable patterns in our genetic data. This is like finding out that every sentence in this ancient language follows the same basic structure. This structure is easily learnable by AI.

To leverage this insight, we built a new model called dnaSORA. This model is a composition of several innovations:

1. It combines two separate models, a generator and a discriminator, into one – an impressive technical feat that reduces architectural complexity and enhances precision.
2. Unlike traditional genetic studies that require thousands of patients to identify disease-causing genes, dnaSORA needs genomes from only a few dozen patients. This is crucial for studying rare genetic diseases with limited available data.
3. dnaSORA uses transfer learning from easily generated synthetic data to make correct predictions on real genomic data, further resolving the training data problem.

These innovations have enabled dnaSORA to isolate disease-causing aspects of the genome at 0.3 megabase (Mb) resolution. For comparison, pharmaceutical companies typically design new medicines at a resolution in the hundreds of megabases, which results in the failure of two out of three medications. Considering that developing a single medicine typically costs about $3 billion, we estimate that the increased resolution dnaSORA provides can lead to 70% cost savings.

Moreover, dnaSORA is generalizable across approximately 10,000 rare genetic diseases. While we’re excited about this prospect, we acknowledge that human genetic Hawaiian-like experiments performed by other groups have had very poor resolution. We hope that by bringing this experiment out of obscurity and combining it with powerful frontier AI like dnaSORA, we can facilitate the development of new CRISPR medications and significantly scale our understanding of the human genome as a first language.

## Introduction

The human genome is a dataset that humans do not read as a first language. In contrast, we naturally comprehend natural language datasets like English. Recently, multiple AI models have gained the capacity to read natural language as first languages (Radford *et al*., 2018, ^1^) However, no AI models have been developed capable of reading the human genome with the same innate understanding as we do with spoken languages.

Beyond insights on the origins of life, reading the human genome as a first language has importance for health. A strand of the human genome is composed of approximately three billion nucleotides, each of which can be considered a word or token. Misrepresentation of *≥*10,000 of these tokens results in ∼10,000 genetic diseases causing a variety of maladies in ∼800 million humans (∼10% of humans) (Verma and Puri, 2021, ^2^).

These 10,000 tokens present an opportunity to attain the first tranche of words for reading the human genome as a first language. This is because these tokens cause easily readable phenotypes in humans enabling straightforward label assignments. What is needed for these assignments is a data interpreter where various phenotypes output identical data with contextual class labels which in this case are simply token positions.

The Hawaiian experiment (also known as Bulk Segregant Analysis) is an approach that is little known outside of the fundamental research sciences. It takes advantage of extreme natural variation (Michelmore and Kesseli 1991, Anderson and Rockman 2022, ^3,4^) by multiplexing genomes with phenotype causing token misrepresentations such that there is progressively strong selection of small nucleotide polymorphisms (SNPs) from one parent at the genomic location causing the phenotype. This results in a species-invariant point cloud of ∼5 points/kB that is roughly Gaussian (Thompson *et al*., 2015, Anderson and Rockman 2022, Minevich *et al*., 2014 ^5,4,6^). Importantly, this Gaussian shape is identical regardless of the phenotype; however the *mu* of the Gaussian centers at the misrepresented token thus revealing its positional class label. The Hawaiian experiment therefore is a satisfactory data interpreter.

The Hawaiian experiment combines ideas from linkage mapping and GWAS but has poor genomic resolution (Shen and Messer 2022, ^7^). A typical workflow involves localizing to a large search space (megabase-sized, Mb) within which one manually inspects for tokens that are misrepresented relative to a standard healthy reference. Compared to competing approaches (Tam *et al*., 2019, ^8^), an important and distinguishing quality is that the span region of the large search space is nearly always correct because the physics of the point cloud are underpinned by the natural laws of meiosis and linkage (Shen and Messer 2022, ^7^). However, identifying misrepresented tokens remains an unmet challenge because of the large search space (Velásquez *et al*., 2017, ^9^).

The Hawaiian experiment has demonstrated remarkable success across a wide range of organisms, from plants to mammals (Minevich *et al*., 2014, Sun and Schneeberger, 2015, Van Leeuwen *et al*., 2012, Obholzer *et al*., 2012, Leshchiner *et al*., 2012, ^6, 10, 11, 12, 13^). However, its relative obscurity in basic science means that it has not yet been conducted in humans. We’ve determined that the vocabulary size of the point cloud for human Hawaiian experiments is at least ten million points. To capture this representation, we chose the diffusion transformer (DiT) as our model of choice. DiTs are primarily employed in image (Dosovitskiy *et al*., 2020, ^14^) and video (Bertasius *et al*., 2021, ^15^) spaces and have shown remarkable scaling. For instance, the video model SORA by OpenAI (Brooks *et al*., 2024, ^16^) is thought to carry the largest representation of any model in existence. Inspired by this representational power, we’ve named our model dnaSORA.

dnaSORA was initially designed to output synthetic Hawaiian genomes for the downstream training of a larger Large Genome Model (LGM) that would discern the positions of misrepresented tokens. However, we realized that for the model to be able to generate such data, it must have an internal representation of the real genome representation the downstream LGM would be leveraging. This propelled us to investigate the DiT not only as a genome generator, but as a genome classifier as well. Such a unified architecture was recently found to be competitive with traditional classifiers in image space (Mukhopadhyay *et al*., 2023,^17^), including zero-shot classification of Stable Diffusion (Li *et al*., 2023, ^18^). This architecture is radically simplified and accordingly reduces training time. We extend this work to DNA space here in *C. elegans* Hawaiian data.

Our goal is to demonstrate how to cost-efficiently obtain at scale the positional labels of a tranche of 10,000 tokens in humans. We note that 1) these 10,000 tokens are the mechanisms causing rare genetic diseases. They thus are the missing information for wide-spread scaling of CRISPR medications (Parums, 2024, ^19^). 2) The Hawaiian experiment is expandable to any phenotype of the sexual diploid, thus the library of potential tokens is well beyond 10,000.

## Related work

The recent rapid innovation of AI models in image and natural language spaces has been paralleled with innovation in protein and nucleic acid spaces (Dalla-Torre *et al*., 2024, Abramson, *et al*., 2024, ^20,21^). Similar to natural language space, the leading models in DNA space are based on cloze tests (i.e., BERT) or autoregressive (i.e., Claude). These include The Nucleotide Transformer (Dalla-Torre *et al*., 2024, ^20^), DNABert2 (Zhou *et al*., 2023, ^22^), and DNAGPT (Zhang *et al*., 2023, ^23^). These models pre-train on thousands of genomes to gain insights on key features of the DNA code in an unsupervised fashion (Dalla-Torre *et al*., 2024, ^20^).

Recently, a new model explored diffusion as a means to generate DNA regulatory regions such as promoters (Sarkar *et al*., 2024, ^24^). Diffusion models are unexplored in DNA space beyond this work.

## Results

### Universality of the Hawaiian experiment

Hawaiian experiments take advantage of extreme natural variation to generate point clouds. For instance, natural genetic differences in *C. elegans* ecotypes from Bristol, England versus Hawaii, USA (hence the name Hawaiian experiment) are used to generate point clouds (Thompson *et al*., 2015, Anderson and Rockman 2022, ^5,4^). In admixed individuals from these regions, point clouds transform from a uniform distribution to a Gaussian-like feature if a misrepresented token originates from only one of the regions (Figure 1). The Gaussian-like feature is due to meiosis, linkage, and is made visible by multiplexing of genomes (Anderson and Rockman 2022, ^4^). Gaussians have a mean or *mu* representing the center of the distribution. The *mu* represents a search space for identifying misrepresented tokens (Figure 2). This approach has successfully identified hundreds of independent, unrelated phenotypes across species, from nematodes (Minevich *et al*., 2014, ^6^), plants (Sun and Schneeberger, 2015, ^10^), arthropods (Van Leeuwen *et al*., 2012, ^11^), zebrafish (Obholzer *et al*., 2012, ^12^) to mammals (Leshchiner *et al*., 2012, ^13^).

**Figure 1:**
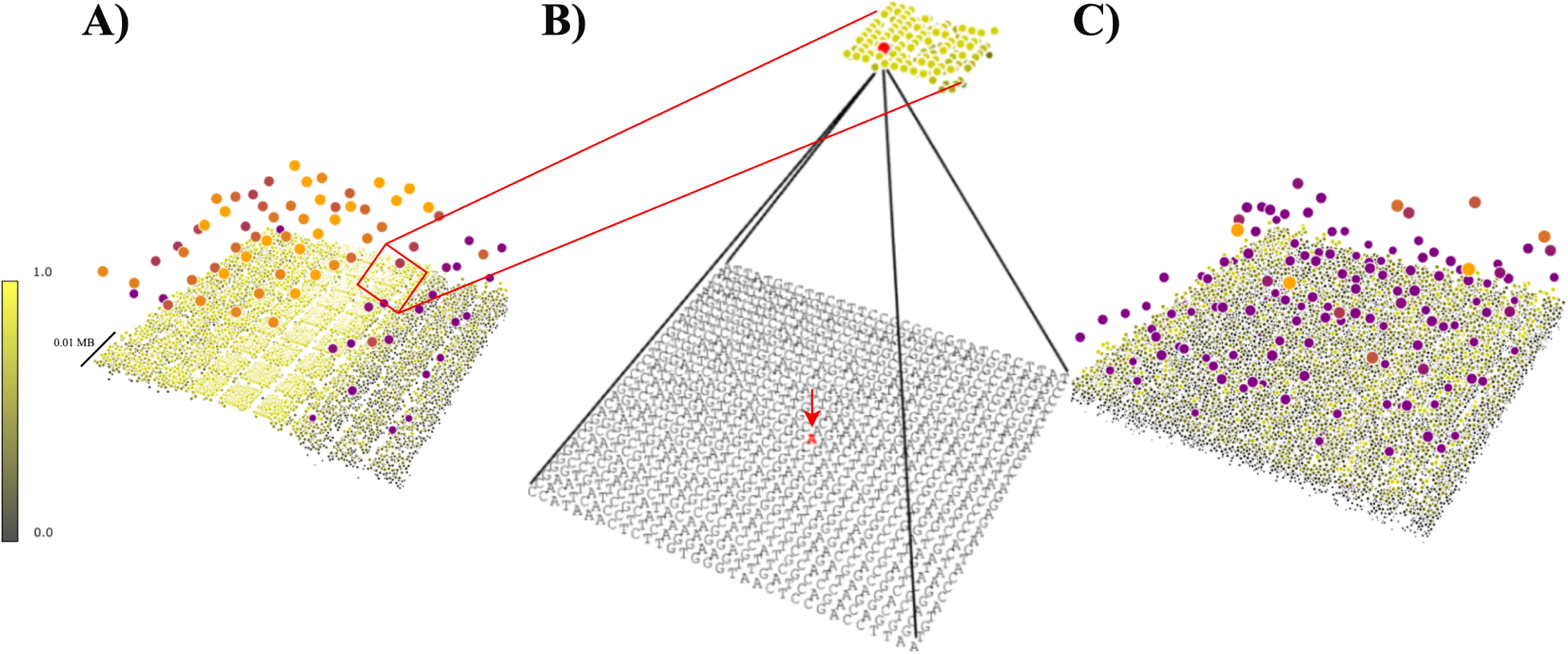
Brooklyn plot of Hawaiian experiment point clouds. A) Representative *C. elegans* chromosome harboring a misrepresented token (i.e., mutation). Bottom plane is a dense point cloud representing local neighborhoods of DNA tokens (i.e., nucleotides). Intensifying yellow hues highlight abundance of tokens from the ecotype the misrepresented token is from. Top plane is a sparse point cloud summarizing the dense point cloud. Intensifying orange hues signify a predominance of one ecotype. B) Zoom of the local neighborhood with a red arrow pointing to a misrepresented token. While the misrepresented token itself is not directly represented in the point cloud, its position is inferred from the characteristics of surrounding tokens. C) Representative chromosome that does not harbor the misrepresented token. Note reduced yellow and orange hues compared to A. Each point symbolizes approximately 1024 tokens. A single chromosome is visualized as a point cloud of roughly 16,000 points, representing about 16 million tokens. See Supplemental Results for rational for naming plot ‘Brookyln’.

**Figure 2:**
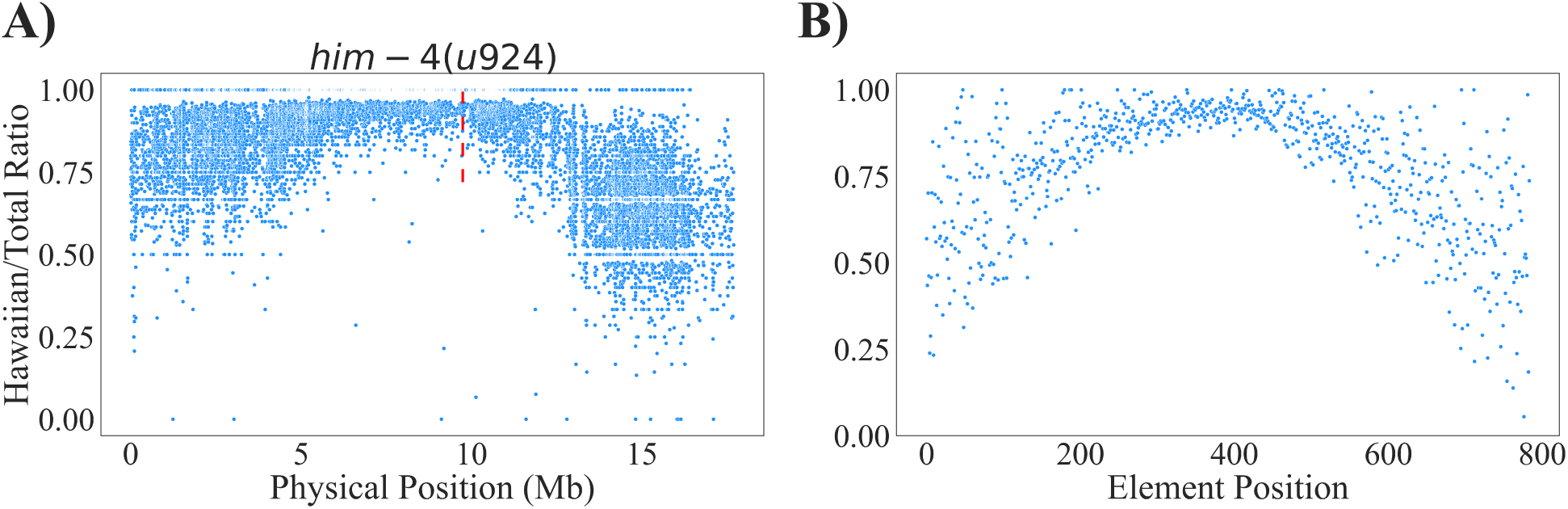
Gaussian feature of real data and model-produced synthetic data A) 2D chromosome point cloud from Hawaiian experiment generated *C. elegans* with a mutation in the hemicentin gene. The mutation, *him-4(u924)*, is a misrepresented token at position 9,742,980 (red line). It causes severe extracellular matrix defects that disorganize the gonad, reducing fertility. Human ortholog is HMCN1. The Gaussian-like feature of the point cloud has a *mu* identifying the location of this token position. B) Representative synthetic data (SD) point cloud generated by the model (max value == 1.2, cutoff == 1). Note similarity of bell-curve in A and B.

### Value of increased token resolution

Experiments that carry some hallmarks of the Hawaiian experiment have been performed on humans (Chi *et al*., 2019, Cullina *et al*., 2023, ^25,26^). These lacked key protocols such as whole genomes, pooling and multiplexing, and were characterized by very poor resolution often in dozens to hundreds of megabases making it challenging to assign misrepresented tokens. Consequently, other methods are more common, though the difficulty with these approaches is they are unlikely to provide correct search spaces (Tam *et al*., 2019, ^8^), To highlight the value of the Hawaiian experiment we explored in a clean room model how point cloud resolution impacts costs in genetic research and therapeutic development by comparing two scenarios: a basic researcher (Agent A) who had studied model organisms, and a large group of independently acting healthcare pharmaceutical companies (Agent B) who had developed CRISPR-based medicines. Both agents investigated 100 distinct genetic targets in 100Mb genomic spaces, where natural error of the DNA replication machinery resulted in ∼75 misrepresented tokens (i.e., *de novo* mutations, background noise) in every newborn human (Kaplanis *et al*., 2023, ^27^). This amounted to two misrepresented tokens per 100Mb alongside each phenotype-causing token.

At 100 Mb resolution, Agent A spent $300,000 total ($3,000 per target) conducting rescue experiments, while Agent B spent $300 billion (B) total ($3B per target) to develop CRISPR medicines, as each had to test three candidates (one correct, two background) per target. Improving resolution to 10 Mb reduced background noise by 90%, requiring only 1.2 tests per target, resulting in costs of $120,000 for Agent A and $120B for Agent B. Further improvement to 1 Mb resolution reduced background noise by 99%, requiring 1.02 tests per target, with final costs of $102,000 for Agent A and $102B for Agent B.

For Agent B, the improvement from 100 Mb to 1 Mb resolution results in a cost savings of approximately $200B or 70%, primarily attributed to the significant decrease in false positives requiring testing due to background noise elimination. See Supplementary Results for baseline assumptions and more details of the clean room model.

### Solving the genome scarcity problem

Despite the Hawaiian experiment having been performed in many organisms, these data are rare. There is no public repository we know of that has such data. We had access to less than 50 Gaussian-like feature containing Hawaiian genomes which is insufficient to train a classifier. We therefore looked at other domains for inspiration. In the image generative AI space, models such as Midjourney (Midjourney, ^28^) and Stable Diffusion (Esser *et al*., 2024, ^29^) are diffusion transformers trained on images including those of the face portraits of real people. These models subsequently generate synthetic face portraits that are so aligned to the data distribution of real faces that humans cannot distinguish synthetic face portraits from real face portraits (whichfaceisreal.com, ^30^). Yet the synthetic faces have variance enough for us to perceive them as unique individuals rather than clones. We wondered if a similar generative model could be created for genomes. In this case, can a model generate synthetic Hawaiian genomes that are near indistinguishable from real Hawaiian genomes, but have variance to make each genome unique? This would solve the data scarcity of Hawaiian experiment genomes as we would have near unlimited data for training a classifier.

### The Unified DiT

While the approach of a synthetic genome generating DiT is new, the idea of using synthetic data for training has been demonstrated elsewhere (Le *et al*., 2017, Zhao *et al*., 2022, Huijben *et al*., 2024, Trinh *et al*., 2024, Zhang *et al*., 2023, ^31,32,33,34,23^). These studies separate synthetic data generation from classification. Indeed the primary use of diffusion transformers such as Stable Diffusion is to generate synthetic data. We reasoned that for these models to generate facial portraits within the distribution of the input data, there must be an internal latent representation of faces that the model is leveraging. Plus this latent space may be a source of representation similar to if not richer than a classifier trained on the outputs of the representation. Several recent studies have explored such internal latent representations and demonstrated that their frozen state can be used for classification (Li *et al*., 2023, ^18^). We therefore investigated whether a DiT trained on Hawaiian experiment genomes can be used as a generator and discriminator (Figure 3).

**Figure 3:**
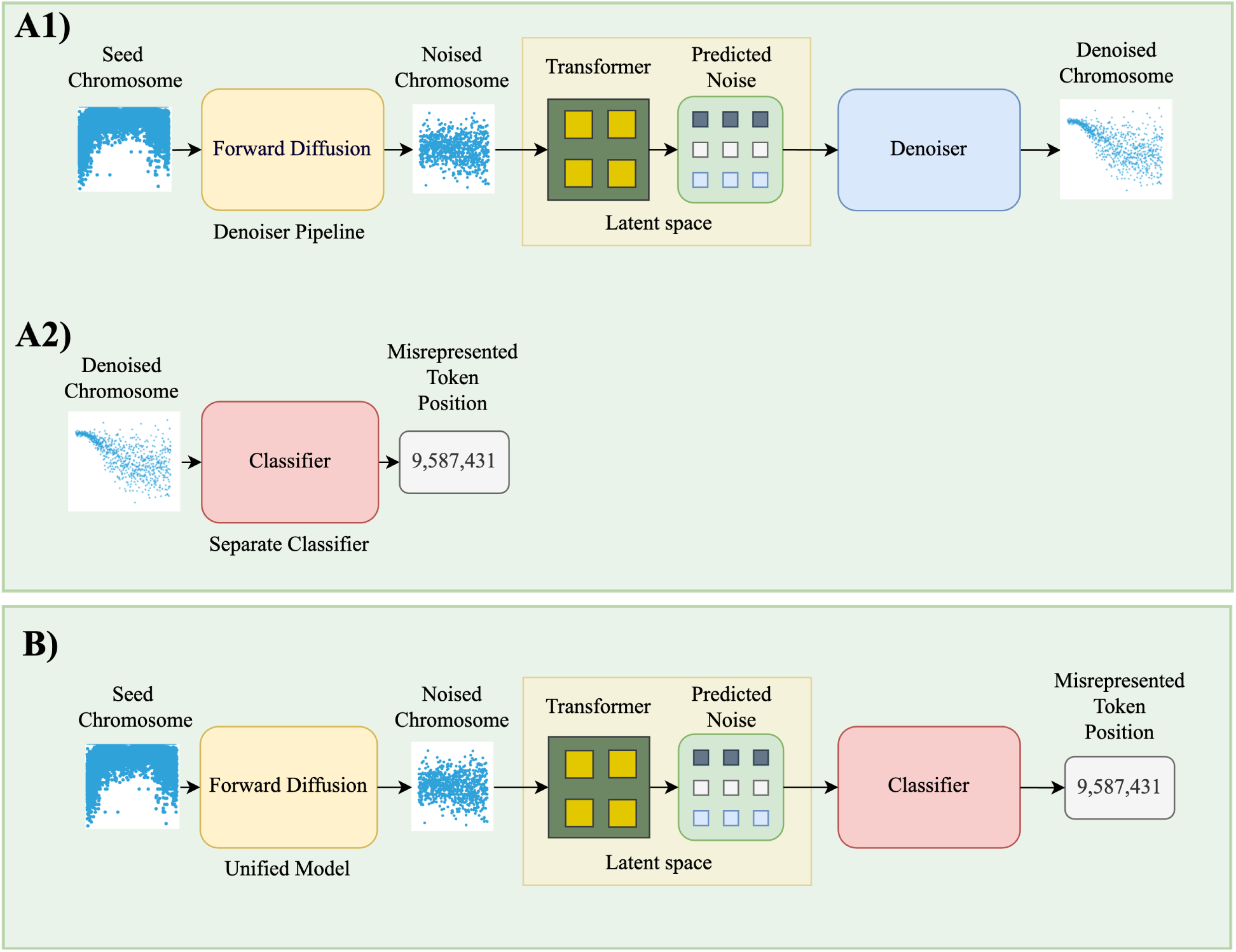
Unified diffusion transformer. Goal of the model is to take in chromosome point clouds as input and output the position of the misrepresented token. A) Non-unified model is a synthetic chromosome point cloud generator. A1) In a conventional approach, this synthetic data is deployed to train a separate classifier (A2) for assigning positions to misrepresented tokens. B) The frozen latent space (see Figure 4) of the generator is leveraged for classification in the unified model.

### Converting a DiT to a Unified DiT

A traditional DiT workflow for classification is composed of a diffuser, transformer, and denoiser which generate synthetic data (SD) for training a separate classifier (Figure 3 A1-A2). To convert the DiT for discrimination, we froze both the DiT neural network and its timestep and class label inputs (Figure 4). We removed the denoiser since it’s use is for creating SD, which was no longer needed as we worked with the transformers frozen latent representation (Figure 4). The model’s classification capability was implemented by adding a linear-layer based head, an attention-based head (Mukhopadhyay *et al*., 2023, ^17^) or a recent unique architecture for zero-shot classification (Li *et al*., 2023, ^18^) we refer to as a zero-shot head.

**Figure 4:**
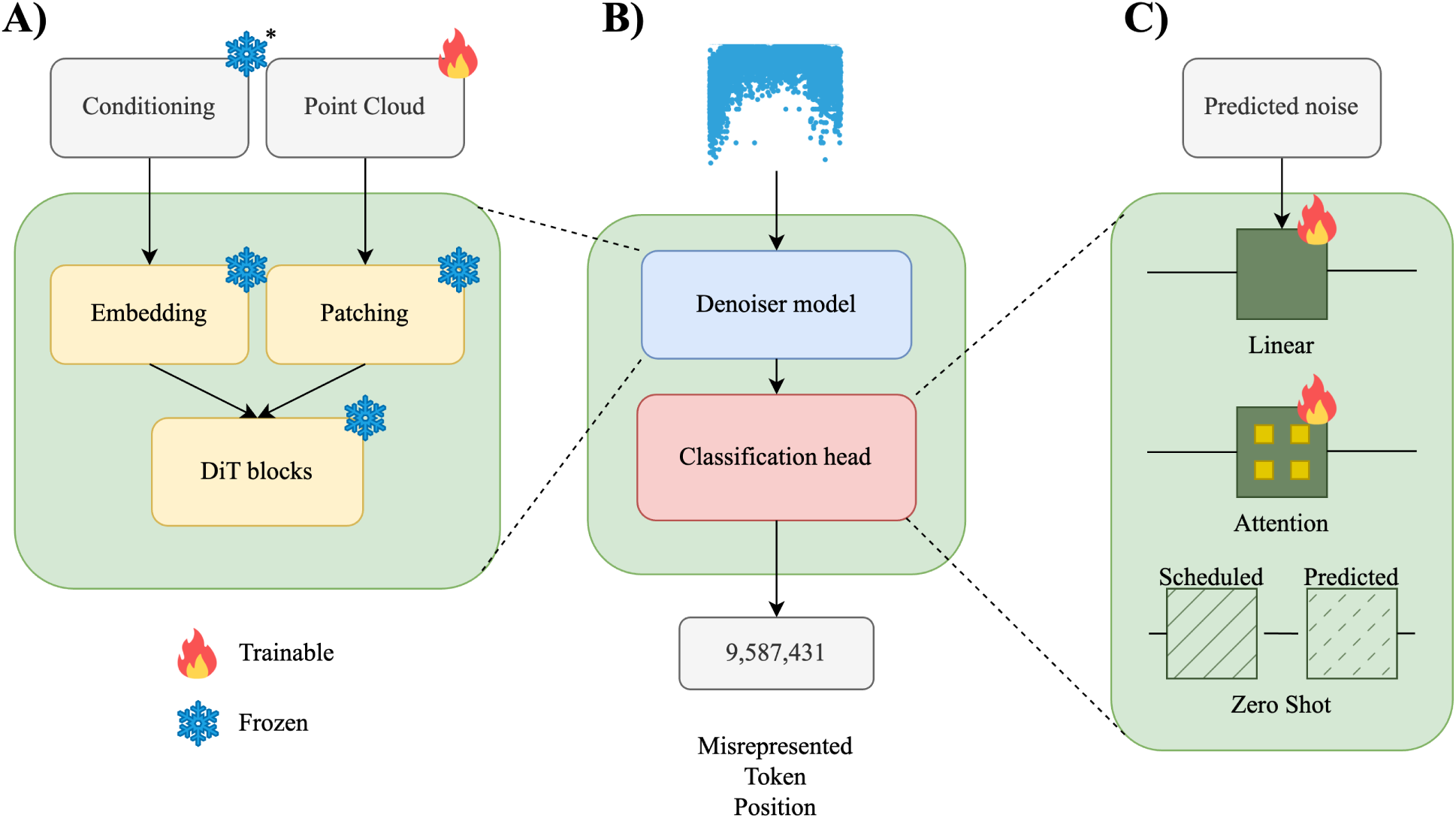
Frozen latent representation. A) The unified model upon training has weights frozen. * indicates aspects of the model that were frozen. B) A classification head leverages the frozen latent representation for discriminator tasks. C) Linear, Attention, and Zero-shot heads were used for classification. * Zero-shot classifier required modification of frozen protocol.

### Types of data used

To build this unified model, we employed several data types. Real data (RD) are Hawaiian experiment C. elegans chromosomes. We had access to 35 of these, three of which have experimental validation of the misrepresented token using the mu from Hawaiian experiments (Figure 2 A). Mock data (MD) are point clouds from several tunable functions (Supplementary Figure S3, Figure S4, Figure S5, Figure S6) to emulate the complexity of RD. For all work presented here, a simple Gaussian function was sufficient. MD is further delineated to sparse MD and dense MD (Supplementary Figure S3). Sparse MD are 784 element point clouds used for rapid training of the models. Dense MD are 16,384 element point clouds (similar to RD) and were used as a stand-in for RD for model accuracy validation. The final data type is Synthetic data (SD) (Figure 2 B). This is the output of the DiT. As we are interested in the internal representation of the DiT, we used SD as a human-readable ‘printout’ to evolve this representation.

### Patch size

Unlike LLMs which employ text tokens, DiT’s commonly employ patches (Brooks *et al*., 2024, ^16^). To input Hawaiian chromosomes into the DiT, we first experimented with parsing them into micropatches as is commonly done in image and video models (Brooks *et al*., 2024, ^16^). The advantage of this approach is that it lends itself well to massive GPU multiprocessing. SD generated with this approach required reassembly of patches into continuous chromosomes. These chromosomes featured RD-like Guassian-like features. However they lacked contiguity between patches. We believe the generative process introduced interpatch variance, a phenomenon also seen in other domains. While there are solutions to this problem, we decided to take advantage of the tractable size of *C. elegans* chromosomes to introduce entire point clouds (i.e., macropatches) into the DiT. Initial experiments demonstrated that a denoising model trained on dense MD could generate SD similar to RD. However, due to slow training performance, we opted to train a reduced DiT model of sparse MD and integrate it into a unified classifier that predicts *mu* for full-length chromosomes. While computationally more intensive, our approach of using contiguous sequences produced contiguous SD exhibiting Gaussian features characteristic of RD.

### Querying for Inference

Querying the model at inference proved interesting because as humans do not understand genomic data as a first language, we cannot query with a prompt that doesn’t make sense to us like ‘AATGCCT’. This contrasts LLMs where one queries with natural language text (i.e., “I’m going shopping”). For Sora, grids of empty patches were employed such that the depth of the grid determined the number of frames of the generated video (Brooks *et al*., 2024, ^16^). The grid size was variable, allowing for video resolution to be specified. Our model is point cloud based, so similar to grids, our model takes at inference patches of point clouds that are variable. We were able to input sparse MD and have the model yield a similar element-sized point cloud. Increasing the size of the point cloud input (i.e., from sparse MD to dense MD or RD) resulted in similar element-sized point cloud output.

Inference for assigning new tokens involves generating new RD which requires pooling of several dozen samples from phenotypic admixed individuals (plants, animals or people) from two to three whole genome sequencing experiments (see Methods).

### Ground truth data

To ascertain phenotype causing misrepresented tokens, our loss function maximized on learning *mu*. Three of the RD; *vab-3(ot266)*, *him-4(u924)*, and *mec-1(u925)* had *mu*’s that previously were used to assign misrepresented tokens experimentally (see phenotypes in Methods). Validation that the misrepresented tokens were causally responsible for phenotypes was 1) observance of nucleotide divergence only in animals with the phenotype, 2) reversion of the phenotype to wild type upon injection of phenotypic animals with DNA fragments containing the correct nucleotide. Thus these three RDs are gold standard ground truths.

The remaining RD do not have this experimental backing. However they have the Gaussian-like feature. We therefore used LOESS curves which are commonly used to approximate the position of misrepresented tokens (William G., 2000, Minevich *et al*., 2014, ^35, 6^).

### Experiments performed

We performed comparisons of performance between the conventional DiT+classifier (referred to henceforth as the conventional model) versus the unified DiT (henceforth referred to as the unified model). Both models were trained on sparse MD. It’s important to clarify that the conventional model comprised two separate models: 1) a sparse MD trained DiT outputting SD, and 2) a classifier trained on this SD (Figure 3 A1-A2). In contrast, the sparse MD trained unified model was a single, integrated model (Figure 3 B).

We measured the performance of the unified model’s classification capabilities using three different approaches:

1. a linear-layer based head, 2) an attention-based head, and 3) a zero-shot head.

**Ground truth based *mu’s:***

- Dense MD had a similar point cloud density as RD and was used as a stand-in for RD for model accuracy validation. Dense MD was generated using functions where we defined the *mu*. We therefore had *a priori* knowledge of *mu’s* for dense MD. We refer to this as dense MD *mu*.
- Our three ground truth RD *vab-3(ot266)*, *him-4(u924)*, and *mec-1(u925)* had *mu* values that were used to assign misrepresented tokens experimentally. Since misrepresented tokens are not elements in our point clouds, we programmatically found the closest point to them, and refer to this point as RD *mu* (Figure 1). The element distance of the misrepresented token to RD *mu* for *vab-3(ot266)*, *him-4(u924)*, and *mec-1(u925)* was 1, 753, and 114 tokens (nucleotides) respectively, so negligible.

**LOESS based *mu’s:***

- The LOESS method is a common approach for discerning misrepresented token positions using Hawaiian point clouds (Minevich *et al*., 2014, ^6^). We applied this approach for model accuracy validation and comparison to the literature.
- We referred to LOESS derived from MD data as dense MD LOESS *mu.* Since we had prior knowledge of dense MD *mu* values, we were able to empirically compare the accuracy of the LOESS approach with synthetic data.
- We referred to LOESS derived from RD as RD LOESS *mu*. Most of our available RD did not have experimentally validated ground truths (see Methods). Therefore, RD LOESS *mu* served as our best approximation to misrepresented token positions. We were able to batch all RD (with and without experimental validation) to derive aggregate data on accuracy.

In summary, we had two measurements for synthetic and real world ground truth data: <u>dense</u> <u>MD *mu*</u> and <u>RD *mu*</u>. And two measurements for LOESS data <u>dense MD LOESS *mu*</u>, and <u>RD LOESS *mu*</u>.

### Model performance

We first describe the conventional model compared to the unified model with a linear-layer based head (Table 1). We measured the likelihood of identifying the correct search space (defined as Accuracy). Data was aggregated, so 50 MD and 35 RD were examined (Table 1).

- The conventional model discerned dense MD LOESS *mu* with 93.85% accuracy (unified model: 94.26%), RD LOESS *mu* with 84.96% accuracy (unified model: 83.69%), dense MD *mu* with 99.04% accuracy (unified model: 98.82%), and RD *mu* with 93.10% accuracy (unified model: 96.57%). Measures for the unified model with additional classification heads follow the same trend and are included in Table 1.
- The consistently reduced scores associated with LOESS measurements empirically highlight the inaccuracy of this commonly used method. Our most important measure is RD *mu*. Here the high scores across experiments demonstrate effective transfer learning of synthetic to real distributions. We note the unified model was 3.47% more accurate than the conventional model.
- We next examined the conventional model and unified model in more detail (Table 2). Rather than reporting on the accuracy of the search space with aggregate scores, we focus on individual RD point clouds for *vab-3(ot266)*, *him-4(u924)*, and *mec-1(u925)*. These RD have *mu* values based on misrepresented token positions that have been experimentally validated. Our primary measure here is Delta (in megabases) to define how physically close our implementations were to predicting the *mu* position directly flanking the misrepresented token (i.e., RD *mu*).
- No other points are more proximal than RD *mu* to the misrepresented token. Therefore, RD *mu* is the theoretical maximum resolution of a point cloud based approach for identifying the position of misrepresented tokens.
- We found that our top three performers were the unified model with the linear-layer based head (Delta

**Table 1.**
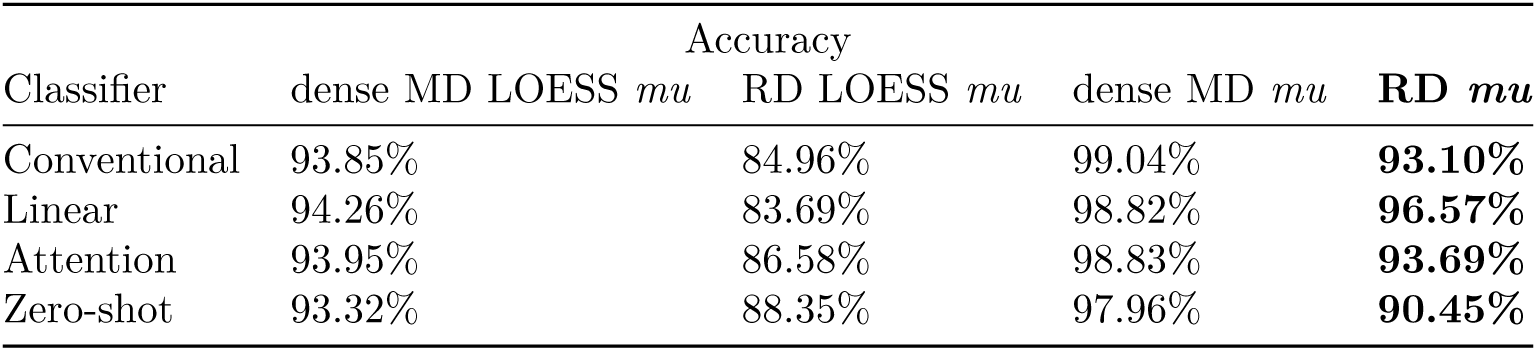

| Classifier | Accuracy |  |  |  |
| --- | --- | --- | --- | --- |
|  | dense MD LOESS *mu* | RD LOESS *mu* | dense MD *mu* | **RD *mu*** |
| Conventional | 93.85% | 84.96% | 99.04% | **93.10%** |
| Linear | 94.26% | 83.69% | 98.82% | **96.57%** |
| Attention | 93.95% | 86.58% | 98.83% | **93.69%** |
| Zero-shot | 93.32% | 88.35% | 97.96% | **90.45%** |

**Table 2.**

| Classifier | Allele | Chrom | RD *mu* | Predicted *mu* | **Delta (Megabase)** |
| --- | --- | --- | --- | --- | --- |
| Linear-layer head | *vab-3(ot266)* | 6 | 10,517,648 | 10,220,326 | **0.30** |
| Zero-shot head | *mec-1(u925)* | 5 | 6,962,458 | 7,260,425 | **0.30** |
| Linear-layer head | *him-4(u924)* | 6 | 9,742,866 | 9,264,843 | **0.48** |
| Conventional model | *mec-1(u925)* | 5 | 6,962,458 | 7,727,019 | **0.76** |
| Conventional model | *vab-3(ot266)* | 6 | 10,517,648 | 11,349,103 | **0.83** |
| Linear-layer head | *mec-1(u925)* | 5 | 6,962,458 | 7,902,164 | **0.94** |
| Attention head | *mec-1(u925)* | 5 | 6,962,458 | 8,065,577 | **1.10** |
| Attention head | *vab-3(ot266)* | 6 | 10,517,648 | 9,234,955 | **1.28** |
| Attention head | *him-4(u924)* | 6 | 9,742,866 | 8,084,289 | **1.66** |
| Zero-shot head | *vab-3(ot266)* | 6 | 10,517,648 | 8,356,084 | **2.16** |
| Conventional model | *him-4(u924)* | 6 | 9,742,866 | 7,475,837 | **2.27** |
| Zero-shot head | *him-4(u924)* | 6 | 9,742,866 | 5,954,007 | **3.79** |

== 0.3 Mb), the zero-shot head (Delta == 0.3 Mb), and again the linear-layer based head (Delta == 0.48 Mb). Curiously, models performed dynamically depending on the input data. This suggests the complexities distinguishing each RD uniquely interacted with each model’s internal representation.

- We next determined if the size of the search space of the unified model was competitive with the literature (Table 3). What distinguishes the Hawaiian experiment compared to competing methods (Tam *et al*., 2019, ^8^) is the search space it provides nearly always correctly contains the misrepresented token because the physics of point clouds are based on the natural laws of meiosis and linkage (Shen and Messer 2022, ^7^). However identifying misrepresented tokens in a large search space can be exceedingly difficult.
- We found in recent *C. elegans* literature that LOESS-like methods were commonly used, which provided search spaces ranging from 3 Mb to 13.8 Mb (Trividi *et al*., 2023, Krauchunas *et al*., 2023, Fung *et*

**Table 3.**

| Experiment | Accuracy |
| --- | --- |
| Low *et al.*, 2019 | 13.8 Mb |
| Fung *et al.*, 2023 | 8 Mb |
| Krauchunas *et al.*, 2023 | 3 Mb |
| Trividi *et al.*, 2023 | 2 Mb |
| Boost Algorithm | 0.82 Mb |
| dnaSORA | 0.3 Mb |

**Table 4.**
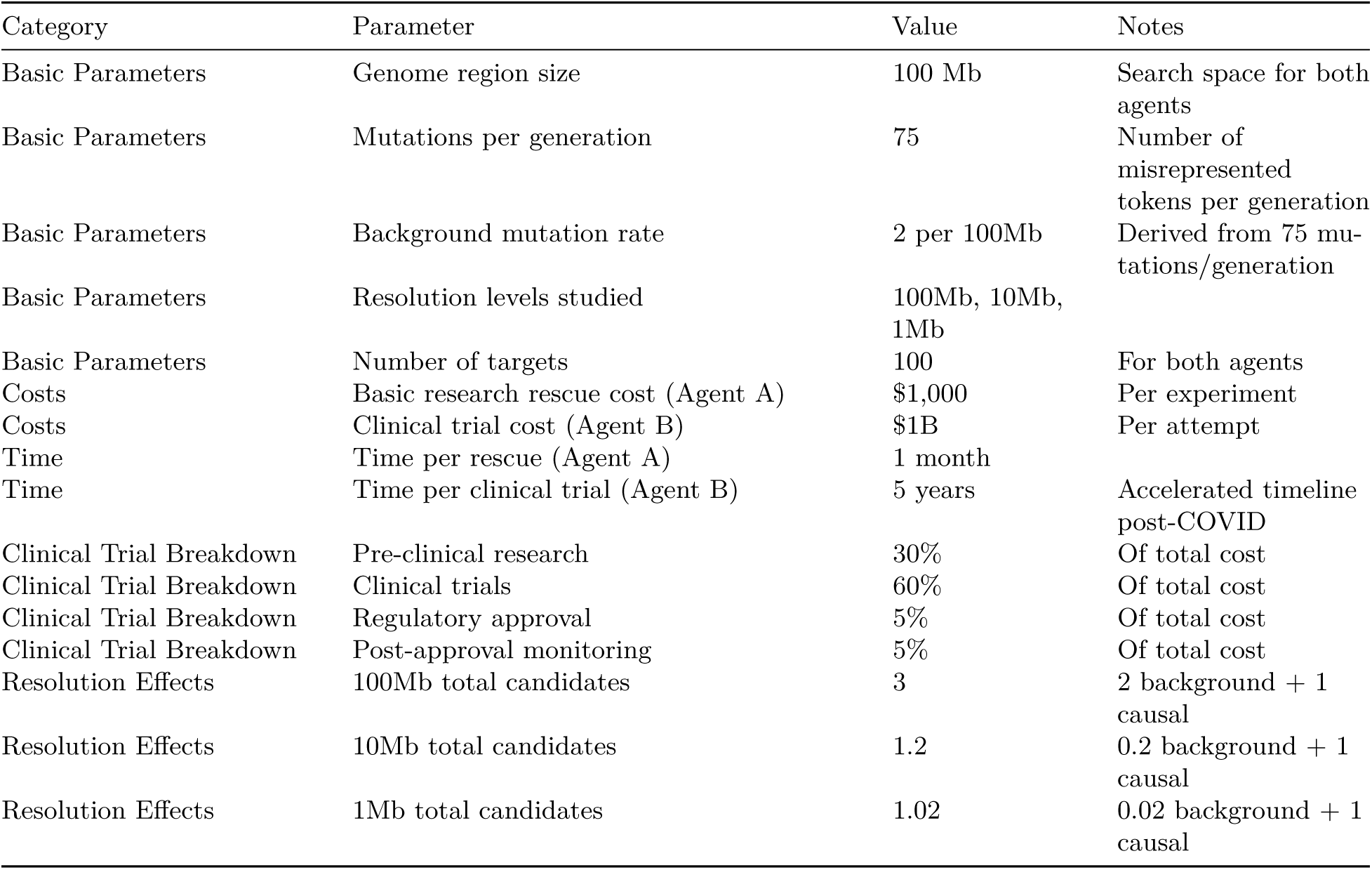

*al*., 2023, Low *et al*., 2019, ^36,37,38,39^). We found use of the Boost algorithm (Sun and Schneeberger, 2015, ^10^) in plant literature. This algorithm compares the observed allele frequencies to the expected allele frequency and in our hands gave a search space of 0.82 Mb for *vab-3(ot266)*. For side-by-side comparison to the Boost algorithm, we inputted *vab-3(ot266)* into our unified model with a linear-layer based head and found its search space to be 0.3 Mb.

Best intervals from misrepresented token reported in other studies using LOESS or similar methodology. dnaSORA and Boost algorithm tested on *vab-3(ot266)*.

## Limitations

We have observed in a random sample of 2000 *C. elegans* genes that 63% feature one or more points. We also observed that despite a ∼30x increase in whole genome size in humans relative to *C. elegans*, the density of point clouds is consistent between the species (∼1000-5000 points/Mb). Human genes are ∼0.03 Mb on average (Fuch *et al*., 2014, ^40^).

*C. elegans* genes, on the other hand, are typically *≤*0.01 Mb. As most of these genes harbor points, we believe the theoretical maximum resolution of our approach is ∼0.01 Mb. Therefore, there is room to improve our approach by a further ∼30x.

An open scientific question is whether every token in the 3B token human genome is assignable to a phenotype. Previous labeling of large chunks of the genome as ‘junk DNA’ has been abandoned as important regulatory regions are found in these intergenic regions (Bernstein *et al*., 2012, ^41^). The trajectory of this research suggests that all 3B tokens indeed have function and thus phenotype labels. This is supported by increased discovery of extremely nuanced phenotypes (Fung *et al*., 2023, ^42^). A key question is which approach will best assign all these tokens? dnaSORA is a supervised approach we propose as potentially appropriate for this task. However, unsupervised approaches such as the Nucleotide Transformer (Dalla-Torre *et al*., 2024, ^20^) may also be able to contribute to this endeavor. Unsupervised language models are able to cluster genomic regions in ways that align with known biological functions (e.g., clustering DNA sequences that we know are promoters based on prior laboratory experiments).

However, there are clusters for which we do not yet understand the biological significance. These could represent either noise in the data or assignments that are beyond our current understanding of the genomic code. This is not necessarily a limitation of Large Genome Models but rather a reflection of the current state of human knowledge. Because of this, we envision supervised models like dnaSORA to be complementary to unsupervised models for progressively assigning labels to genomic elements.

This first iteration of dnaSORA treats tokens and Gaussian-like features as discrete objects. This is the case despite the Hawaiian experiment capacity to capture multigenic information (i.e., a single phenotype with several *mu’s* pointing to multiple tokens) (Minevich *et al*., 2014, ^6^). Thus given a 10-mer element space of 1 to 10, were elements 2 and 8 have important connectivity, say a *cis* and *trans* multimer, dnaSORA will not reveal this information. We believe scaling the library size of Gaussian-like features together with introduction of self-supervised masked discrete diffusion blocks (Sahoo *et al*., 2024, ^43^) to the model will result in the emergence of long distance meaning. This has parallels to the scaling from GPT-1 to GPT-3 with natural language.

## Methods

### Real Data source

Whole genome sequences were derived from *C. elegans* mutants. These were generated from mutagenesis (Brenner, 1974, ^44^) of the Bristol, England derived laboratory wildtype N2 ecotype and subsequent cross to the genetically divergent Hawaii derived ecotype CB4856 (Doitsidou *et al*., 2010, Anderson and Rockman 2022, ^45,4^). Point clouds of allelic probabilities were generated from the whole genomes at around 150,000-300,000 points of divergence (i.e., SNPs) between the N2 and CB4856 ecotypes (Anderson and Rockman 2022, ^4^). These were termed Real Data (RD). Meiosis, linkage and positive selection of phenotypic admixed individuals resulted in progressive abundance of N2 derived points precisely proximal to the causative mutation (i.e., misrepresented token) (Doitsidou *et al*., 2010, ^45^). This progressive abundance is the cause of the Gaussian-like feature in the RD point cloud. We select for the *mu* of the feature as it points causally to the misrepresented token. 35 human curated Gaussian-like feature containing chromosomes from whole genome sequences of 16 *C. elegans* strains were used. The misrepresented token position of only three of these chromosomes were known. These were of strains with alleles *vab-3(ot266)*, *him-4(u924)*, and *mec-1(u925)* (generated by O. Hobert and M. Chalfie labs) with misrepresented tokens that previously were validated experimentally with rescue experiments. VAB-3 is an ortholog of human PAX6. It is a homeodomain transcription factor needed for dopamine neuron specification. HIM-4 is ortholog of human hemicentin (HMCN1) and needed for extracellular matrix organization particularly in the gonad for normal fertility. MEC-1 is an EGF and Kunitz domain containing extracellular protein needed for neuronal ensheathment and mechanosensation (Doitsidou *et al*., 2010, Emtage *et al*., 2004, ^45,46^). 20-60 animals were pooled for whole genome sequencing of each strain. Whole genome experiments (on newly pooled animals) were repeated at least twice for redundancy.

### Mock Data generation

To generate MD, we employed a Gaussian probability density function. Sparse MD were 784 length point clouds and dense MD were 16,384 length point clouds. Sparse MD was used for pre-training and dense MD for inference. 120,000 sparse MD was used for pre-training the conventional and unified models.

### Point cloud distillation

For inference, we distilled input data by slicing chromosomes to create phase offsets based on element indexes. For instance, given a point cloud of elements 1 to 10, distilled representations 1, 3, 5, 7, 9 and 2, 4, 6, 8. 10 exist. Distillation resulted in approximately 300-600 representations per chromosome. The final *mu* prediction is derived from the averaged measurements across all representations. Distillation allowed modest GPU resources to efficiently parse high dimensional chromosomal data.

### Base architecture

We had a choice of autoregressive or diffusion architectures to implement our model. We chose diffusion due to literature suggesting efficient representation of massively high dimensional data (Brooks *et al*., 2024, ^16^). These are essential because whole genomes scale immensely across Kingdoms. Our DiT follows recent work favoring transformers for diffusion (Peebles *et al*., 2022, ^47^). Briefly, a forward diffusion process progressively injects noise to input training data according to a scheduler. A transformer exposed to this data learns to predict the information needed to remove the noise added by the scheduler. It leverages this in a reverse denoising process to generate new data within the data representation of the training data (Supplemental Figure S2). Further details, particularly important nuances in pre-training and inference can be found here (Peebles *et al*., 2022, ^47^).

Our model took as input 1D data. We transformed input data to 2D patches using a convolutional layer and projected them into a high-dimensional feature space. We also conditioned the model with class labels *mu*, and a timestep value (0-1000) indicating the noise level from the scheduler. The input data is normalized, then scaled, and shifted using parameters from the conditioning vector, then processed through attention and feed-forward layers with residual connections to produce the final latent representation. This latent representation is learned by transformer blocks with adaptive Layer Normalization Zero (adaLN-Zero) (Peebles *et al*., 2022, ^47^). Each block features multi-head self-attention mechanisms where the conditioning signals modulate layer normalization parameters through learned scale and shift operations.

Querying Sora for inference involves inputting grids of variable sized empty patches (Brooks *et al*., 2024, ^16^). Inference of our model occurs similarly. One notable difference is Sora processes all patches at once with interpatch information to enable continuity across video frames. Our model processes one patch at a time. This ‘macropatch’ is the entire chromosome point cloud and thus inherently continuous. Also note that the shape of the model output is fixed to the input query shape.

### Separate conventional classifier

The separate classifier model is a classifier with standard features including convolutional blocks, batch normalization, max pooling, etc. (O’Shea and Nash, 2015, ^48^).

### Unified classifier

Previous work has shown that DiT latent representations can be frozen and leveraged for classification (Li *et al*., 2023, ^18^). We froze conditioning inputs (class and timestep), all layers including transformer blocks, patching and embedding blocks, in our model (i.e., frozen latent representation), and added classification layers similar to Mukhopadhyay *et al*., (Mukhopadhyay *et al*., 2023,^17^) yielding us our unified model. We explored three classification layers that varied in complexity. 1) A linear head that consisted of a single linear projection (Mukhopadhyay *et al*., 2023,^17^). 2) An attention-based head with a series of transformer blocks with multi-head self-attention (Mukhopadhyay *et al*., 2023,^17^).

1. 3) An alternative architecture we’ve termed the zero-shot head due to previous results (Li *et al*., 2023, ^18^). Unlike the other heads, the zero-shot head unfroze classes for the purpose of predicting *mu* by comparing the range of predicted noise to scheduled noise (Li *et al*., 2023, ^18^).

### Additional information

The models were pre-trained on datasets of sparse MD samples. We observed that the batch size should be equal to or a multiple of the total label count for denoiser and class count for the classifier for optimal load balancing on all labels. Work was done with pyTorch and supplemental libraries, on RTX3090 GPUs (Nvidia, Santa Clara, CA). The reproducible codebase is available at our GitHub.

## Discussion

Recent advancements in Large Language Models have demonstrated that AI can learn natural language as a first language (Radford *et al*., 2018, ^1^). However, no AI exists that reads the human genome as a first language. Towards this goal, we’ve developed dnaSORA, a unified diffusion architecture that leverages Hawaiian experiment phenotype representations for assigning labels to DNA tokens. dnaSORA’s resolution of 0.3 Mb is best in class.

Generating large genome models requires thousands of individuals that each require sequencing (Dalla-Torre *et al*., 2024, ^20^). Therefore such models are expensive and unscalable. In contrast, pre-training dnaSORA is of negligible cost as synthetic data is used to generate its internal representation. Inference requires pooling several dozen individuals for two to three sequencing experiments. Lastly, our projections on human studies suggest enhancement from 100 Mb to 1 Mb resolution will reduce clinical trial failure rates threefold. Collectively, our solution may accelerate CRISPR therapeutic development at orders of magnitude lower cost.

A key insight in this work is that the Hawaiian experiment is an excellent data interpreter. It’s approach, which is relatively little known outside of fundamental research sciences, presents thousands of distinct phenotypes as a singular gaussian-like feature. For instance, a genetic disease that deteriorates lymphatic tissues in the body has the same data representation as a genetic disease that deteriorates brain tissue. dnaSORA learns these representational features for token annotation. The approach is theoretically scalable to the three billion tokens of the human genome.

dnaSORA is a unified diffusion transformer where we excavate its internal latent representation for classification. This novel architecture (Mukhopadhyay *et al*., 2023, ^17^)) is notably distinguished from conventional diffusion and autoregressive architectures. Model excavation is an emerging field that has been used for alignment (Xia *et al*., 2022, ^49^). The authors of Sora suggest this model is a world simulator (Brooks *et al*., 2024, ^16^). To generate realistic videos, its latent space must contain fundamental functions of nature for motion, gravity, and so forth. Excavating this latent space may refine known functions and reveal unknown functions. Similarly, we foresee training of dnaSORA on more complex genetic code, and excavating its latent space to understand the human genome as a first language.

## Supplemental Results

### Value of increased token resolution

We explored how DNA sequencing resolution dramatically impacted costs when searching for misrepresented tokens, by comparing two scenarios: a researcher who had studied model organisms (Agent A) and a large group of independently acting healthcare pharmaceutical companies who had developed patient therapies (Agent B). Using real-world values, we created an idealized clean room model to examine this relationship.

### Assumptions

- Our model operated in 100 Mb genomic spaces. While whole genome sequencing typically has 0.1% to 1% error rates, we assumed perfect accuracy for this analysis. Both agents investigated 100 distinct targets where misrepresented tokens caused phenotypes - for Agent A, these were phenotypes like single neuron migration defects in basic research, while for Agent B, these represented severe neurological diseases requiring therapeutic intervention. Agent A aimed to understand these phenotypes, while Agent B’s goal was to develop CRISPR-based treatments for all 100 conditions.
- Each generation (i.e., newborn baby) introduced approximately 75 misrepresented tokens (i.e., *de novo* mutations) diverged from parents, equating to roughly 2 misrepresented tokens per 100 Mb (Kaplanis *et al*., 2023, ^27^). This background rate of misrepresented tokens applied equally to both agents’ systems.
- In basic research, rescue experiments validate causal relationships between misrepresented tokens and phenotypes. This involves identifying candidate misrepresented tokens, creating corrected DNA fragments with the healthy version of the token, introducing them to gametes, and confirming phenotype correction in offspring. For Agent A, using *C. elegans* at 100 Mb resolution, each rescue experiment cost was approximately $1,000 and took one month.
- The therapeutic equivalent to a rescue experiment was bringing a CRISPR medication to market, which cost approximately $3B over 5-10 years. This broke down into pre-clinical research (30%), clinical trials (60%), regulatory approval (5%), and post-approval monitoring (5%). With 90% ($2.7B) being research-related costs, we conservatively estimated Agent B’s costs at $1B per attempt (Wouters *et al*., 2020, ^50^), with an accelerated five-year timeline reflecting post-COVID efficiencies.
- We assumed Agent B never determined that the misrepresented token is incorrect early on using preclinical screening, which include *in vitro*, *ex vivo*, and other vetting efforts. Cost savings are variable depending on when the screening works, thus difficult to account for. We assume the industry-wide $1B average cost of bringing medicines to market (Wouters *et al*., 2020, ^50^) captures all instances of successful screening attempts.
- Agent B’s approach differed from basic research by targeting somatic cells (like brain ventricular cells) rather than gametes, adhering to ethical guidelines.
- Both agents utilized the Hawaiian experiment methodology, ensuring the misrepresented tokens they seek existed within their 100Mb search spaces.
- While time can be optimized through parallel processing (e.g., batch sequencing, multiple companies), costs remained independent due to the unique nature of each misrepresented token that must be investigated.
- We use fractional values for scaling (i.e., 0.1 clinical trials makes little sense, but 0.1 clinical trials x 100 does). At 100Mb resolution
- Agent A generated small genomic fragments containing the correctly represented tokens for all three candidates (one phenotype-causing token plus two background tokens) and performed the rescue experiment on three groups of animals. The offspring of one of these groups no longer had the phenotype, thus revealed to Agent A the identity of the misrepresented token. The cost per target was $3,000 ($1,000 × 3). For 100 targets, the total cost was $300,000 ($1,000 × 3 × 100; representing 100 correct rescues and 200 incorrect rescues from background noise).
- Agent B had developed three CRISPR medications per target and had put them through clinical trials at a cost of $1B each. Only one - the one that corrected the misrepresented token - would succeed. The total cost per target was $3B. For 100 targets, the total cost was $300B ($1B × 3 × 100; representing 100 correct medications and 200 failed trials from background noise).

At 10Mb resolution

- The background token misrepresentation rate was reduced by a tenth, from 2 per 100Mb to 0.2 per 100Mb. Therefore, Agents A and B assigned 1.2 total misrepresented tokens per target (0.2 background tokens plus 1 phenotype-causing token).
- For Agent A, the cost per target became $1,200 ($1,000 × 1.2, representing one correct rescue plus 0.2 incorrect rescues). For 100 targets, the total cost was $120,000 ($1,000 × 1.2 × 100).
- For Agent B, the cost per target became $1.2B ($1B × 1.2, representing one correct medication plus 0.2 incorrect trials). For 100 targets, the total cost was $120B ($1B × 1.2 × 100).

At 1Mb resolution

- The background token misrepresentation rate was reduced by another factor of 10 from the 10Mb resolution, going from 0.2 per 100Mb to 0.02 per 100Mb. Therefore, Agents A and B assigned 1.02 total misrepresented tokens per target (0.02 background tokens plus 1 phenotype-causing token).
- For Agent A, the cost per target became $1,020 ($1,000 × 1.02, representing one correct rescue plus 0.02 incorrect rescues). For 100 targets, the total cost was $102,000 ($1,000 × 1.02 × 100).
- For Agent B, the cost per target became $1.02B ($1B × 1.02, representing one correct medication plus 0.02 incorrect trials). For 100 targets, the total cost was $102B ($1B × 1.02 × 100).

This shows how improving resolution from 100Mb to 1Mb dramatically reduced the number of false positives (background tokens) that need to be tested, resulting in significant cost savings for both agents.

### The progression shows

- 100Mb: 3 tests needed per target (1 correct + 2 background) | $3B for 100 CRISPR medicines
- 10Mb: 1.2 tests needed per target (1 correct + 0.2 background) | $120B for 100 CRISPR medicines
- 1Mb: 1.02 tests needed per target (1 correct + 0.02 background) | $102B for 100 CRISPR medicines

To summarize for Agent B, which was a group of large healthcare pharmaceutical companies pursuing 100 different targets, the cost at 100 Mb resolution was $300 billion (requiring 3 trials per target), at 10 Mb resolution was

$120 billion (requiring 1.2 trials per target), and at 1 Mb resolution was $102 billion (requiring 1.02 trials per target). While increasing resolution does reduce costs by decreasing the number of background tokens that need to be tested, the relationship isn’t strictly linear with resolution because there is always the constant cost of the one true phenotype-causing token that must be tested. The savings come from the reduction in false positives that must be tested due to background noise.

### Internal Latent Representation Interpretation

- An in-depth examination of the generative process is insightful for understanding our leveraging of the model’s internal latent representation. The denoising model receives complex 1D chromosome point cloud data of probabilities that is a noise-modified data. Within the model is a neural network that receives this data as a 2D transform and adds further dimensions to the vector space. To the human eye, the transformed 2D chromosome is effectively a grayscale image (noise space, which we’ll call *image X*) (Figure S1).
- The forward process of diffusion involved the neural network learning the noise that was added to point clouds from chromosomes (chromosome space). The neural network, given this new *image X* utilizes this knowledge to predict the noise (Figure S1) needed to have *image X* transform from noise space to be a timestep closer to chromosome space. This prediction is then subtracted from grayscale *image X*, completing a denoising cycle. This cycle repeats multiple times over on *image X* and results in a progressively clearer representation of a nascent chromosome point cloud. This chromosome point cloud has the features of the training set sparse MD while remaining wholly unique (similar to synthetic face portraits from Stable Diffusion).
- The chromosome point cloud exists not only as the output SD but also as an internal latent representation within the neural network. SD can be analogized to a physical paper printout of a boat. The internal latent representation of SD is equivalent to the memory space of the boat in the printer’s RAM chip. However, while RAM data is ordered and cohesive over runs, neural representations are comparatively stochastic. Using the unified model, we excavate this representation.
- We introduce the term “Brooklyn plot” to describe Figure 1, distinguishing it from Miami plots and, more notably, Manhattan plots (Chi *et al*., 2019, Cullina *et al*., 2023, ^25,26^). While Manhattan plots are characterized by data points that dramatically spike upwards from a baseline, reminiscent of the iconic New York City skyline, Brooklyn plots offer a different visual metaphor. Much like the relatively flat skyline of Brooklyn, these plots present a more uniform appearance at first glance. However, just as Brooklyn’s neighborhoods boast unique characteristics that may not be immediately apparent, Brooklyn plots reveal subtle yet significant patterns and clusters of data that reward closer examination.

---

**Figure S1:**
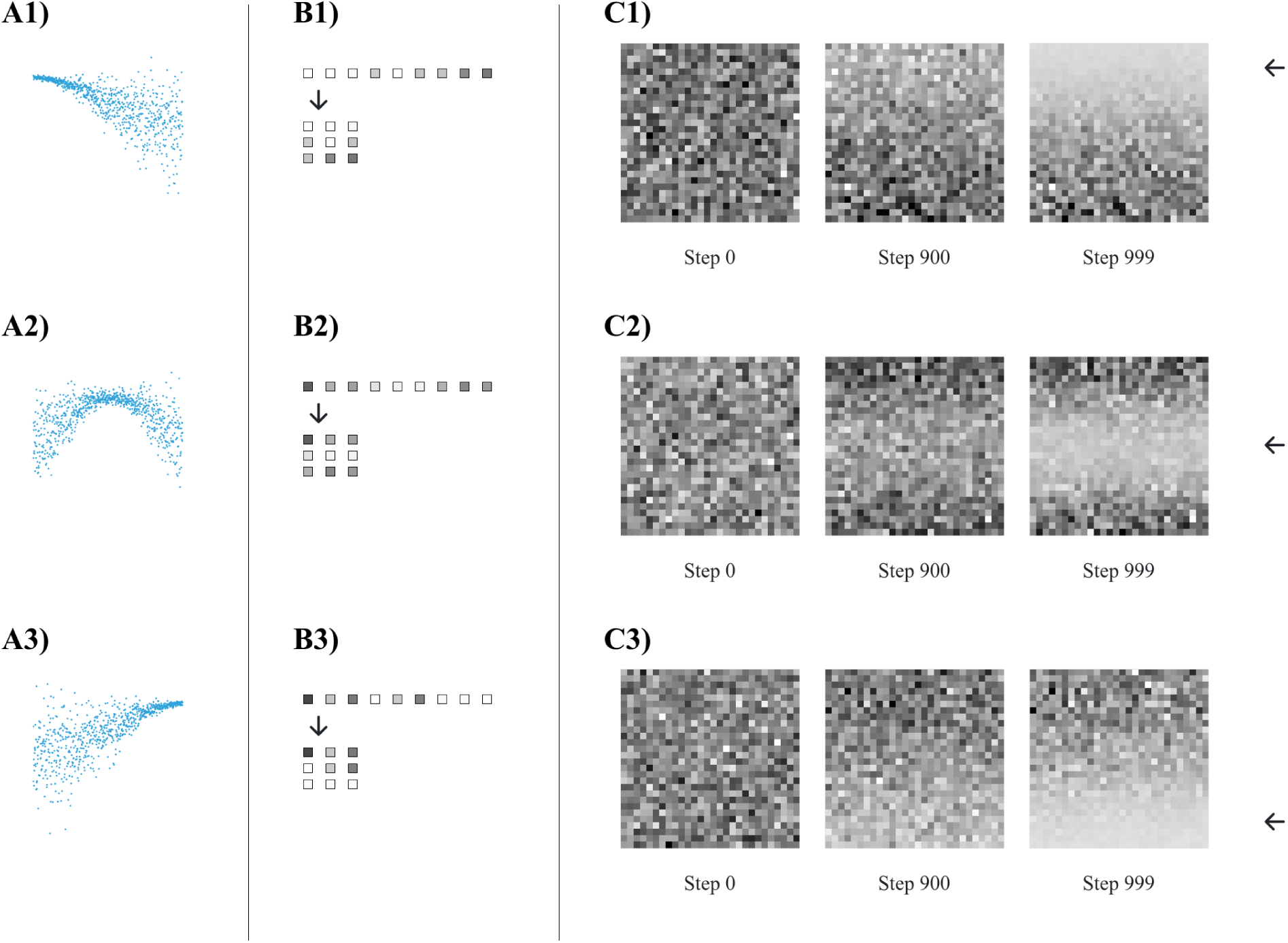
Model interpretability. A) Representative dense MD that are left, middle and right shifted. B) 1D representation of point cloud where each element is shaded according to its Hawaiian ratio (i.e., values closer to 1 are dark; pure Bristol, England derived points are 1, and pure Hawaii derived points are 0). Down pointing arrows indicate transformation of 1D representations to 2D that are learned by the model. C) 2D representations are sequentially denoised according to the scheduler revealing a white band feature. As the *mu* positions are left, middle and right shifted, the position of the nascent white band to be top, middle and right oriented (left pointing arrows). The white band feature is likely an important aspect of the learned latent representation.

**Figure S2:**
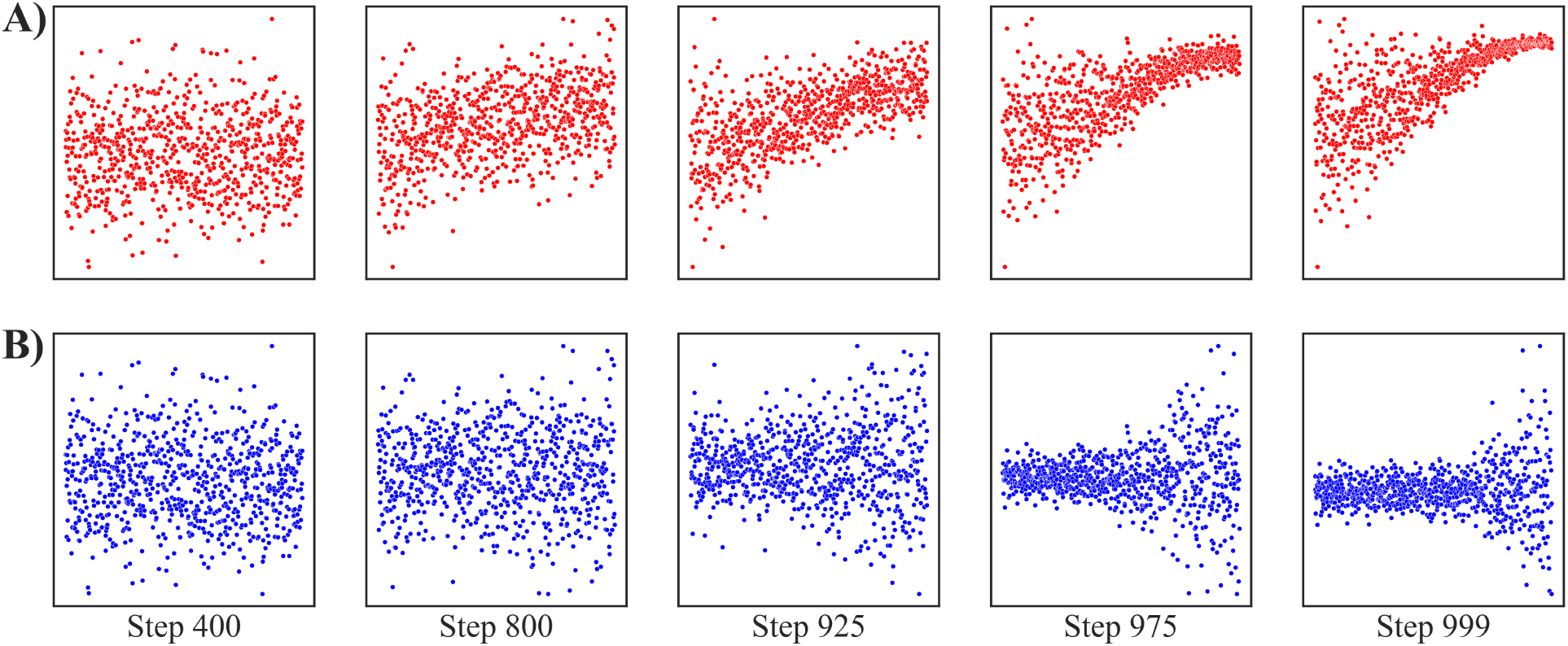
Denoising evolution. A) Progressive sculpting of chromosome point cloud from denoising reverse process. B) Transformer generated predicted noise used to sculpt.

**Figure S3:**
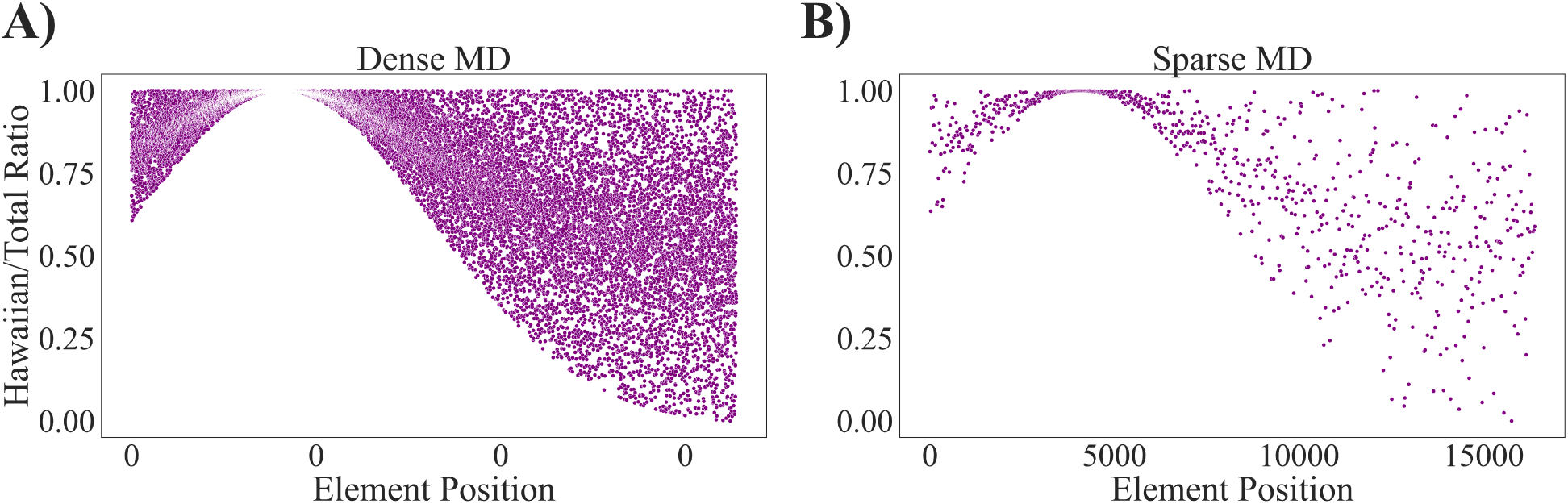
Mock data. A) Dense MD containing 16,384 points and B) sparse MD containing 784 points. Point clouds generated with a Gaussian probability density function.

**Figure S4:**
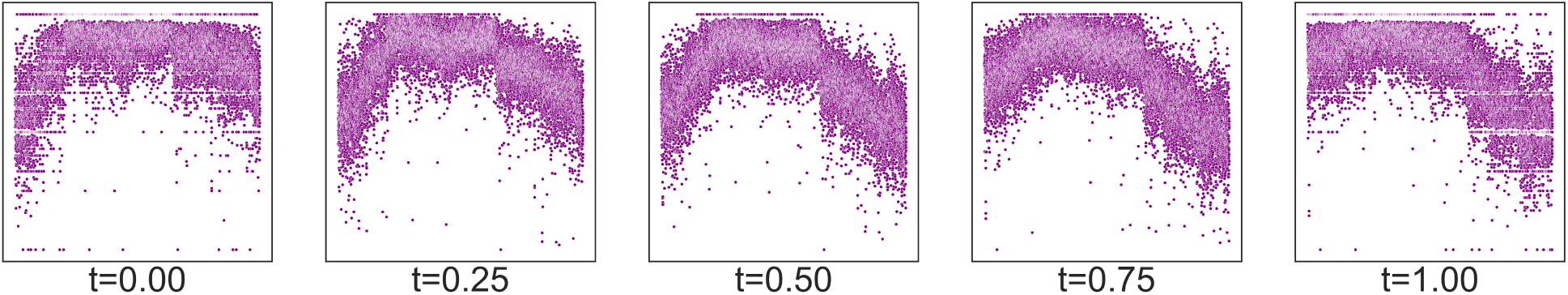
Spherical interpolation captures complexities of RD. Interpolation of point clouds from two RD chromosomes generates new MD that is complex. It represents the strength of interpolation.

**Figure S5:**
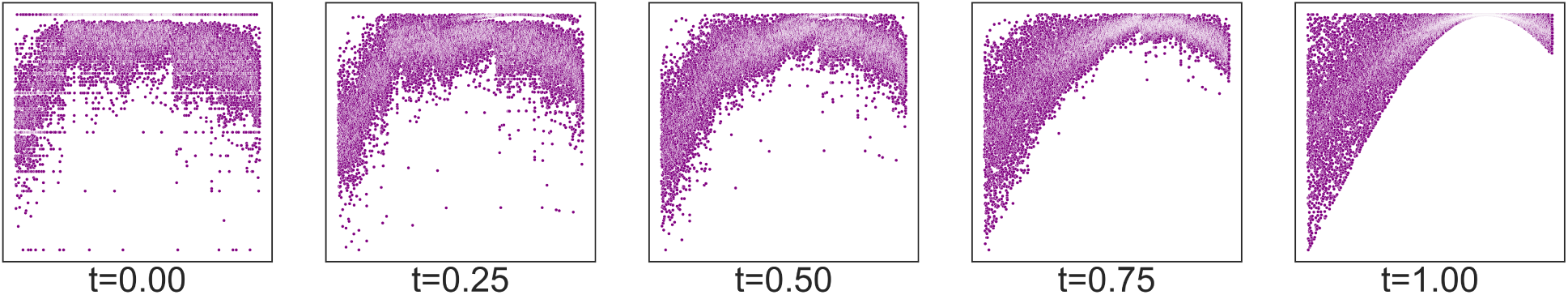
Spherical interpolation for morphing RD. Spherical interpolation of point clouds from complex RD to simple MD. Training data can be generated from desired level of complexity.

**Figure S6:**
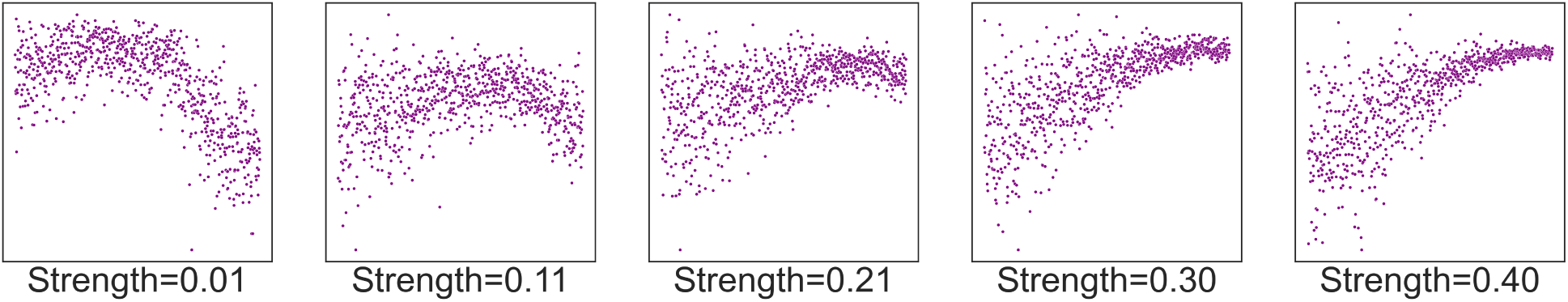
Tuning SD with chr2chr. Condition values such as *mu* for the conventional model are adaptable to explicitly morph SD. We refer to this process as chr2chr and is similar to img2img where an existing image is re-imagined in Stable Diffusion (Rombach *et al*., 2021, ^51^).

